# From solo to collaborative: the global increase in neuroscience authors over two decades

**DOI:** 10.1101/2024.08.05.606580

**Authors:** Mariella Segreti, Ann Paul, Pierpaolo Pani, Aldo Genovesio, Emiliano Brunamonti

## Abstract

The increase in the number of authors per article is a well-documented phenomenon across various academic disciplines. While prior studies have examined this trend in specific fields and countries, they in most cases did not compare the increase in the number of authors between countries. While it has previously shown that the number of authors in neuroscience publications has risen in the G10 countries, no study has yet addressed whether it reflects a global trend in the field. To address this gap, we quantified the global trend in the number of authors in neuroscience publications from 2001 to 2022.

Our findings reveal a consistent increase in authorship across nearly all the countries examined. Italy ranks highest in terms of average authorship per article, while Ukraine ranks the lowest. On the other hand, China shows the largest increase in authorship over the years, followed by Norway and Egypt. South Korea is the only country showing a slight decreasing trend rather than growth. These results contribute to a better understanding of authorship patterns in neuroscience, and can stimulate further investigations on the reasons behind such an increase in terms of socio-economic factors, the need for collaborative efforts in some fields, or, on the negative side, the effect of utilitarian reasons to meet career evaluation criteria.

## 1. Introduction

The increase in the number of authors per paper has been shown across various academic disciplines (Khan et al., 1999; Papatheodorou et al., 2008; Fox et al., 2016). For instance, Fernandes and Monteiro (2016) examined the evolution in the number of authors of computer science publications, considering over 200,000 article references over a 60-year period, from 1954 to 2014. Their study found a consistent increase in the average number of authors per paper across all decades. Moreover, Jang and colleagues (2016) investigated the increase in the number of authors per paper in Korean science and technology publications from 2000 to 2015, finding that the global trend of authors growth per paper is evident in Korea as well.

However, there is limited literature addressing the increase in the number of authors when considering differences between countries. Indeed, while other aspects such as citations, self-citations (Baccini & Petrovich, 2023) and the number of publications has been investigated globally (Baccini & Petrovich, 2023) and reported on platforms like SCIMAGOJR (https://www.scimagojr.com/), the global trend in the number of authors in neuroscience remains almost unexplored.

A first step was taken in our previous study (Paul et al., 2024), where we analysed the rise in the number of authors in neuroscience publications within the G10 countries. However, no study has yet addressed the global trend in the number of authors and its growth, particularly in neuroscience.

To address this question, our current study aims to investigate the global trend in the number of authors in neuroscience publications. In this way, we aim to better understand the collaborative patterns in neuroscience across countries.

## 2. Methods

We adopted a methodology analogous to our previous study on the increasing number of authors in neuroscience (Paul et al., 2024). The dataset for this analysis comprised all neuroscience publications from 2001 to 2022, sourced from 306 neuroscience journals indexed in the Web of Science (WoS, www.webofscience.com).

In this study, we included only countries with a minimum of 450 total publications over the 2001–2022 period. This threshold was selected for two reasons. First, we posited that a smaller number of articles would not provide a reliable average number of authors per article, as a limited dataset is more susceptible to the influence of outliers. Second, many countries with fewer than 450 articles displayed years with zero publications. By applying this threshold, we analysed 49 countries, encompassing a total of 843,231 publications. The results from some of the countries have been already published in our previous article (Paul et al., 2024) and are included for comparison.

Moreover, the number of authors per publication was anomalously high in some cases, with some publications exceeding 400 authors. Possible explanations for this high number have been already discussed in (Paul et al., 2024). Following the methodology of our previous article (Paul et al., 2024), we excluded publications with more than 40 authors.

### 2.1. Data analysis

As in our previous study on this topic, we used a MATLAB script generated in that previous work to identify the country name from the address of the corresponding author. The number of authors was determined by counting the semicolons (;) in the author list and adding one, as the last author is followed by a dot and not by a semicolon.

We then constructed a matrix containing the number of authors per paper, the country of the corresponding author, and the year of publication. This data was used to calculate the average number of authors per year from 2001 to 2022. To assess changes in authorship patterns over time, we employed the delta measure, comparing differences between two sub-periods (2001–2011 and 2012–2022).

To statistically test whether the number of authors differed significantly between countries and across periods, we performed a two-way mixed ANOVA (49 × 2), with Country as the between-subjects factor and Sub-period as the within-subjects factor. All statistical analyses were performed using both MATLAB R2021 (www.mathworks.com) and SPSS (https://www.ibm.com/spss).

## 3. Results

Figure 1 provides a global overview of the average number of authors per country in neuroscience.

**Figure 1.**
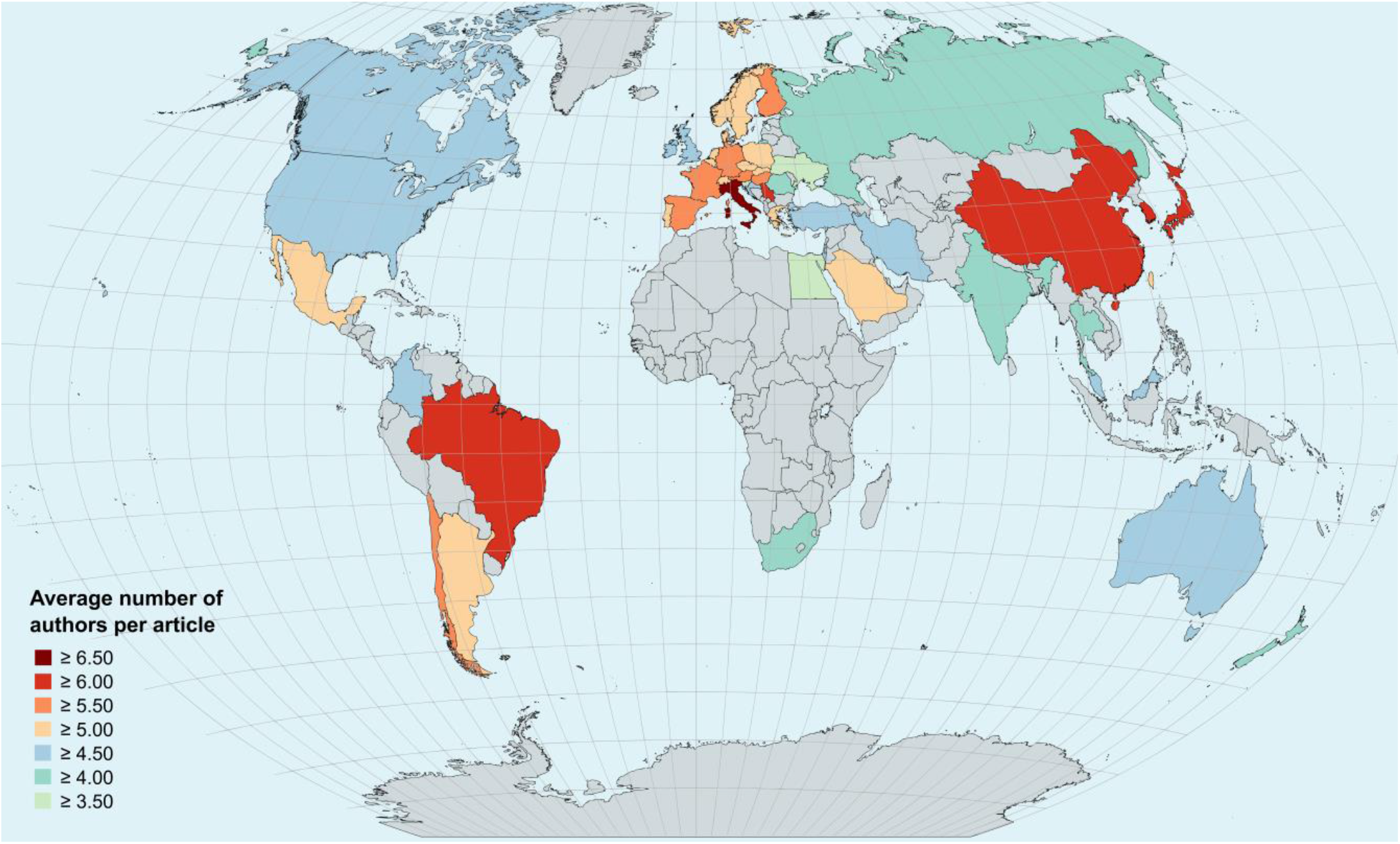
The average number of authors per country in the field of neuroscience. Only countries with a minimum of 450 published articles were included in this analysis, while other countries are coloured in grey. The map was created using MapChart (https://www.mapchart.net/).

The figure shows that Italy has the highest average number of authors, with 6.5 authors per article, followed by South Korea, China, Serbia, Japan, and Brazil, each with more than 6 authors per article.

While Canada is the G10 country with the lowest number of authors, as shown by Paul et al. (2024), we observed that other countries, including nations like Egypt and Ukraine, have even lower numbers. The ranking of the 49 countries is reported in detail in Supplementary Table 1, which also reports a quantification of the growing number of authors per publication calculated as a delta between the 2012-2022 and 2001-2011 decades. The magnitude of this delta for each county is graphically represented in Figure 2.

**Figure 2.**
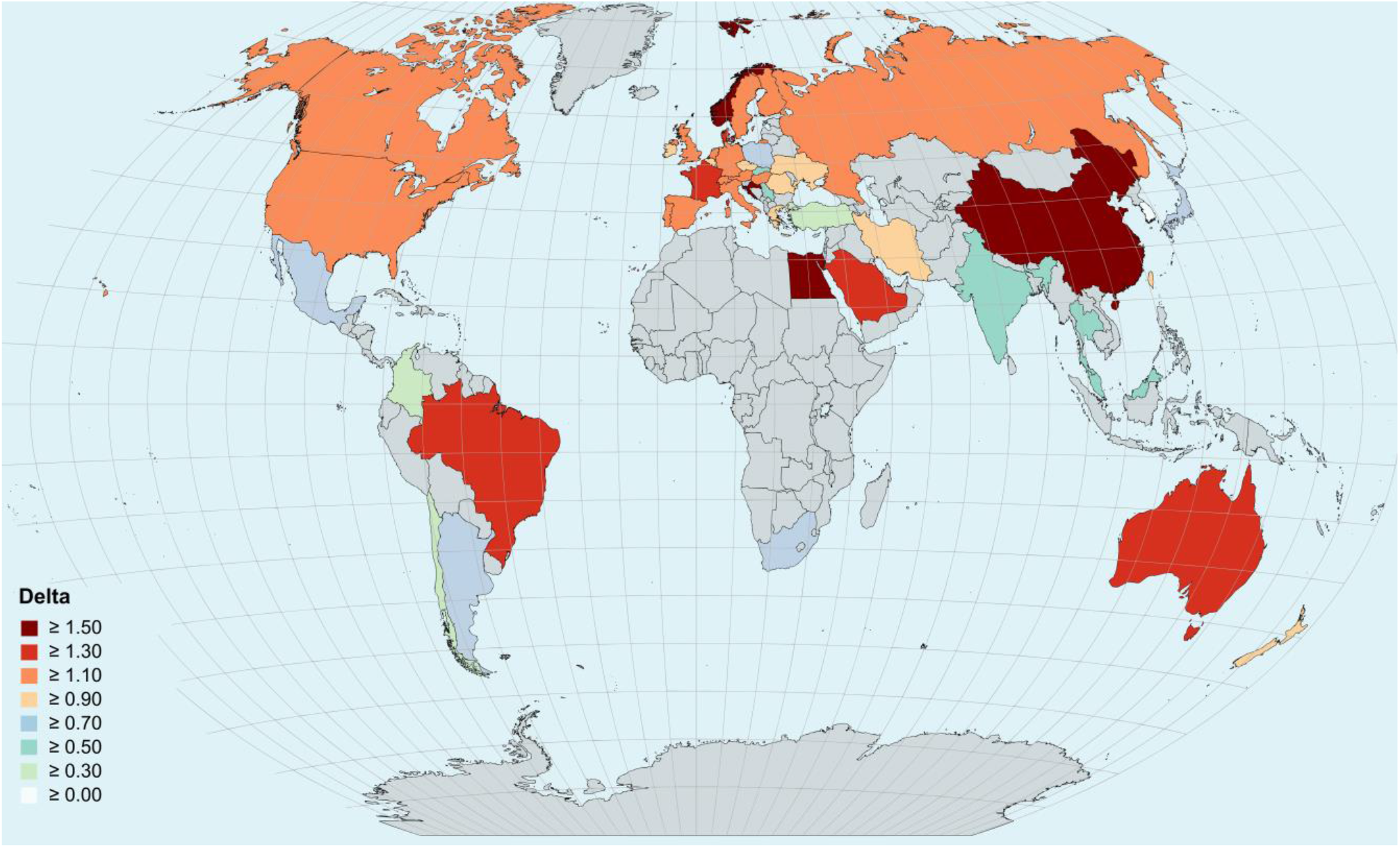
Map displaying the delta values in the field of neuroscience, representing the change between these two sub-periods (2001-2011; 2012-2022), for each country. Only countries with a minimum of 450 published articles were included in this analysis, while the other countries are coloured in grey. The map was created using MapChart (https://www.mapchart.net/).

A two-way mixed ANOVA revealed that the number of publications was significantly influenced by both the Country (F(48,490)=21.24, p < 0.001) and Sub-period factor (F(1,490)=1328.2, p < 0.001), that highlights a significant increase of the average number of authors in the 2011-2022 decade. Additionally, the interaction between the two factors was statistically significant (F(48, 490)=3.21, p < 0.001). China shows the largest delta between the two sub-periods, followed by Norway and Egypt. South Korea is the only country showing a slight decreasing trend rather than growth (Figure 2).

Newman-Keuls post-hoc comparisons between countries indicates that the number of authors of the first-ranking countries (Italy, South Korea, China, Serbia, Japan, Brazil) was significantly higher than the rest of the countries (all Ps < 0.05), while it did not differ between them (all Ps > 0.05). Similarly, the number of authors of the lowest ranking (Romania, Thailand, New Zealand, Russia, India, Slovenia, Egypt, Ukraine) differed significantly from the rest of the countries (all Ps <0.05) while their values were comparable (all Ps >0.05). Country-by-country Newman-Keuls post-hoc comparisons are presented in Supplementary Figure S6. More in-depth details on the growing trends from 2001 to 2022 are illustrated in supplementary figures S1, S2, S3, S4, and S5, depending on the continent (Africa, Oceania, Asia, America, and Europe) each country belongs to.

## 4. Discussion

Examining the increase of authors world-wide we found that the increasing trend of authorship in the field of neuroscience is a global phenomenon. This pattern persists even in countries where the number of publications is relatively low, although the analysis was restricted to countries with at least 450 articles. However, the number of articles is also gradually increasing in countries that are currently underrepresented, such as many in Africa and South America. This is evidenced by data from the SCIMAGOJR site (https://www.scimagojr.com/), which displays the number of publications on Scopus for each country from 1996 to 2023.

Italy stands out as the country with the highest average number of authors per article in the field of neuroscience. On the other hand, regarding the largest growth from the first sub-period (2001-2011) to the second (2012-2022), China exhibited the highest increase in authorship, followed by Norway and Egypt.

The rapid growth in authorship in China can be partially attributed to the “Publish or Perish” culture that is widely prevalent in the country (Tian et al., 2016). This pressure is particularly intense regarding publishing in international journals and collaborating with authors from other countries. Indeed, Fu and colleagues (2011) conducted a bibliometric evaluation of highly cited papers from 1999 to 2009 using the Essential Science Indicators (ESI) database across 22 scientific fields, including neuroscience. They found that 47% of all Chinese ESI papers were international collaborations involving 101 countries. Furthermore, Fu’s analysis revealed that a significant number of the most cited papers were authored by numerous individuals, with some papers having over 200 authors from more than 8 countries. Therefore, the pressure for international publications and the extensive involvement of multiple countries could be significant factors contributing to the substantial growth in the number of authors in Chinese articles.

The increase in authorship in Egypt can possibly be explained, by taking a cue from a non-neuroscientific field. In fact, a study by Farahat (2002) examined authorship patterns in the agricultural sciences in Egypt by analysing all papers published in 19 scientific journals. The findings showed a dominant trend towards multiple authorships, with a steady increase in multi-author papers from 69% in 1960 to 89% in 1980. According to Farahat, a reason for the increasing cooperation is the pressure to collaborate to share the publication costs. As these costs increase, there is a greater incentive for researchers to involve colleagues in sharing the expenses, often resulting in the addition of their names as co-authors, irrespective of their actual contributions to the work.

This rationale presents a plausible explanation, and consequently, it can be anticipated that as the number of publications increases in other nations facing substantial economic challenges, the number of contributing authors will similarly rise. This explanation was overlooked in Paul et al. (2024) but should be expected to account, at least in part, for the increasing trend in authorship in other countries to meet the increasing financial costs associated with the publication process, especially since many journals have moved toward open access, shifting the cost of publication from the readers or university subscriptions to the authors.

However, this rationale carries less weight in economically prosperous countries such as Norway. Nevertheless, Nylenna and colleagues (2014) investigated practices among Norwegian researchers by administering a questionnaire to researchers and PhD students at Oslo University Hospital and the University of Oslo (2700 individuals). They found that 36% of respondents had experienced pressure to include authors in their papers who had not made significant contributions. This could be another of the reasons for the significant growth in the number of authors in such settings.

Finally, the considerable growth in the number of authors per article in Italy can be accounted for, at least in part, by the inclusion of honorary authors as a response to the stringent academic evaluation processes employed within the country, particularly in the context of higher education and research institutions (de Santis Puzzonia et al., 2018). Indeed, in Italy, advancement to ranks such as associate or full professorship is contingent upon meeting specific benchmarks that include the number of publications, citations, and h-index values over a defined period. These criteria are part of a broader framework managed by the Agency for the Evaluation of the University and Research System (ANVUR), which is responsible for setting and recalibrating these thresholds across various academic fields and ranks. The impact of such evaluation criteria on publication practices is profound, because it incentivizes researchers to increase their publication output, often through collaborative efforts, to meet the required thresholds.

Therefore, the reasons behind the worldwide increase in the number of authors per article are not only linked to the growing complexity of scientific subjects (Weeks et al., 2004; Abt et al., 2007) but also to factors related to the economies of countries, peer pressure, and an academic system that relies predominantly on quantitative indicators. This convergence of factors highlights a significant ethical dilemma that has been thoroughly examined in our previous publication (Paul et al., 2024) and that we re-examine only briefly here.

The ethical concern arises from the prevailing “Publish or Perish” culture within academia, which employs publication metrics as critical criteria for career advancement. In countries like Italy, these metrics often can lead to a proliferation of multi-authorship where not all listed contributors have played a meaningful role in the research. This ethical issue highlights the need for normalisation by the number of authors to assign a fair score to the individual scientist, as proposed for the H index (Schreiber, 2008; Galam, 2011; Zerem, 2017). The strong impact of the “Publish or Perish” culture is evident not only in the increasing number of authors but also in the rise of self-citations in certain countries. In fact, Baccini and Petrovich (2023) compared self-citation rates across 50 countries between 1996 and 2019. They found that, while self-citations generally decreased over time in most countries, 12 countries (Colombia, Egypt, Indonesia, Iran, Italy, Malaysia, Pakistan, Romania, Russia, Saudi Arabia, Thailand, and Ukraine) showed the opposite trend.

Our findings on the high number of authors in Italy and the rapidly increasing trend in Egypt align with the high number of self-citations. In contrast, in our work, the other 10 countries identified by the study of Baccini and Petrovich (2023) are not ranked among the countries with the highest number of authors. This difference may suggest that researchers in different countries respond to diverse incentives and institutional pressures regarding citations and publication metrics.

While we identified factors related to the economies of countries, peer pressure, international collaboration, and an academic system that relies on quantitative indicators, further studies addressing the system of incentives and country-specific socio-economic factors are needed to understand these differences. Moreover, while in this study we extended our initial research (Paul et al., 2024) to a global scale, we believe it is equally important to conduct the analysis using a country-centred approach. This would involve examining the increase in the number of authors by geographic areas, specific universities, or even departments to understand the effect of local policies. We have limited our study to the field of neuroscience, but a more comprehensive study across all fields is needed, as some of our results may be specific to this field.

## Data availability

The entire dataset used for this analysis is available at: Paul, A. (2024). Neuroscience publications (2000-2022) [Data set]. Zenodo. https://doi.org/10.5281/zenodo.11445060

## CRediT authorship contribution statement

**MS:** Conceptualization, Data curation, Formal analysis, Writing – original draft, Writing – review & editing, Methodology, Software. **AP:** Conceptualization, Data curation, Formal analysis, Writing – original draft, Writing – review & editing, Methodology, Software. **AG:** Conceptualization, Formal analysis, Supervision, Validation, Writing – original draft, Writing – review & editing. **PP:** Conceptualization, Writing – review & editing. **EB:** Conceptualization, Formal analysis, Supervision, Validation, Writing – original draft, Writing – review & editing.

