## Supplementary Material for "From solo to collaborative: the global increase in neuroscience authors over two decades"

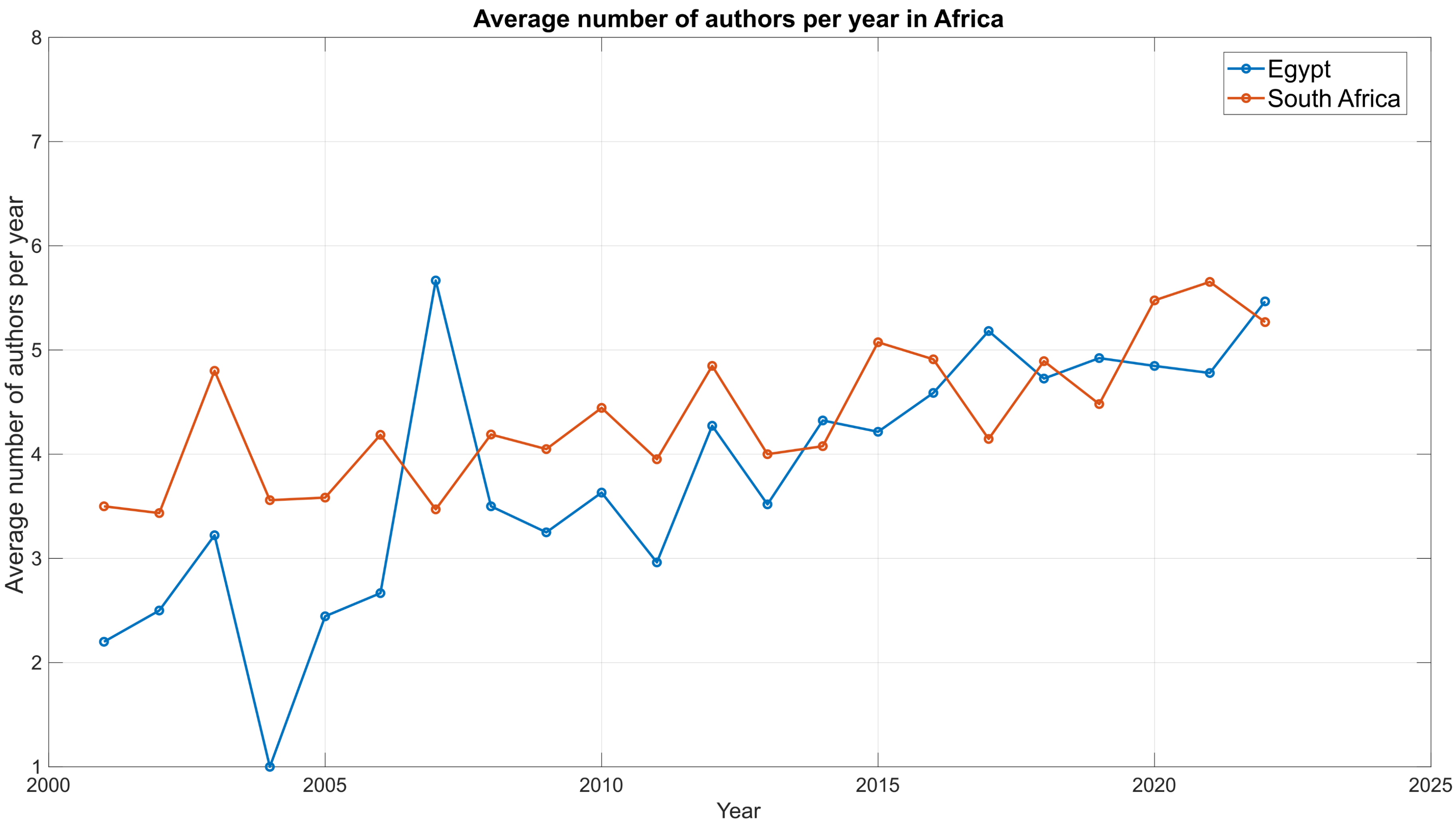


**Figure S1**. Temporal trend of authorship in the field of Neuroscience for the African target countries from 2001 to 2022.

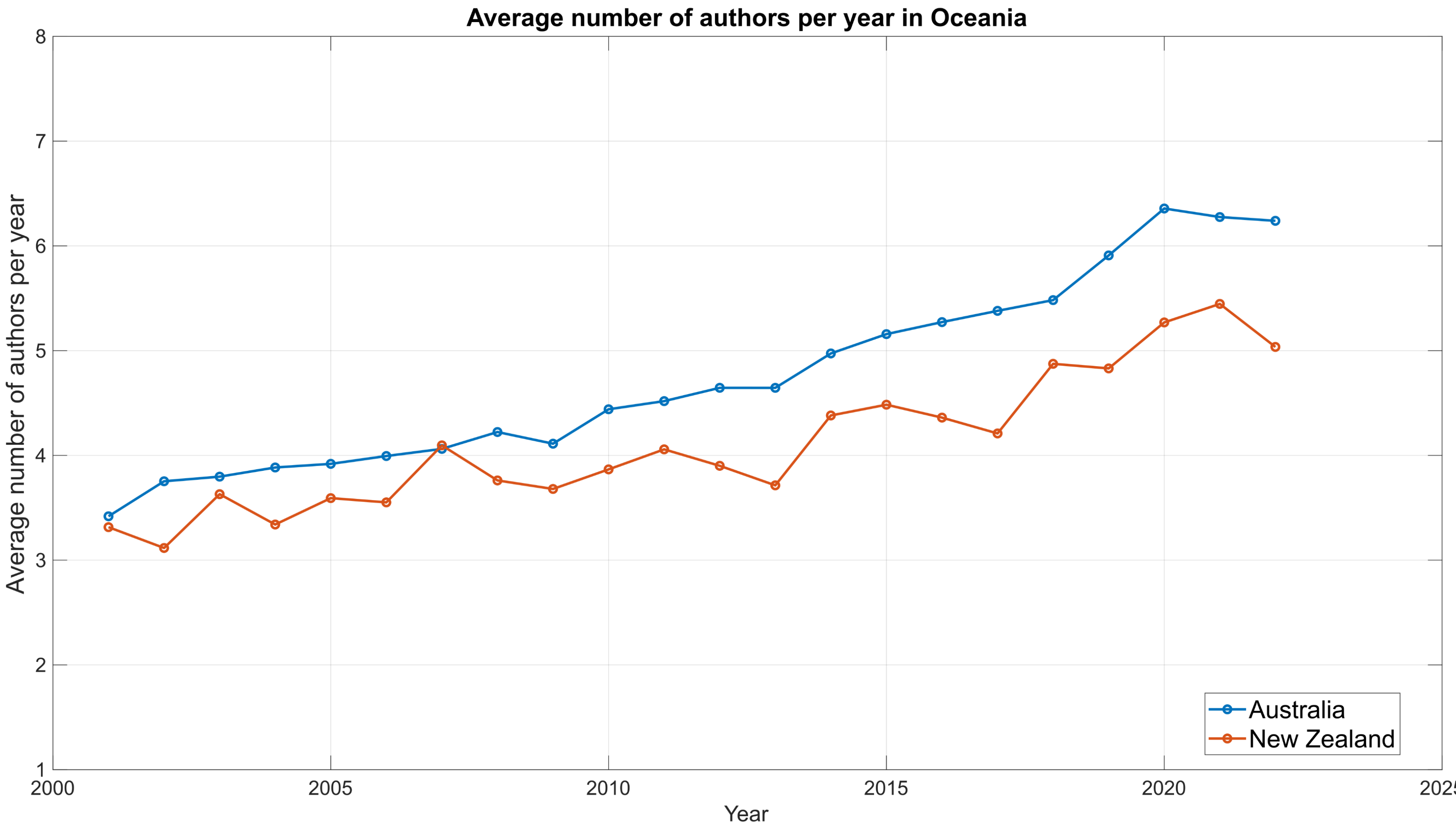


**Figure S2**. Temporal trend of authorship in the field of Neuroscience for the Oceanic target countries from 2001 to 2022.

**a**

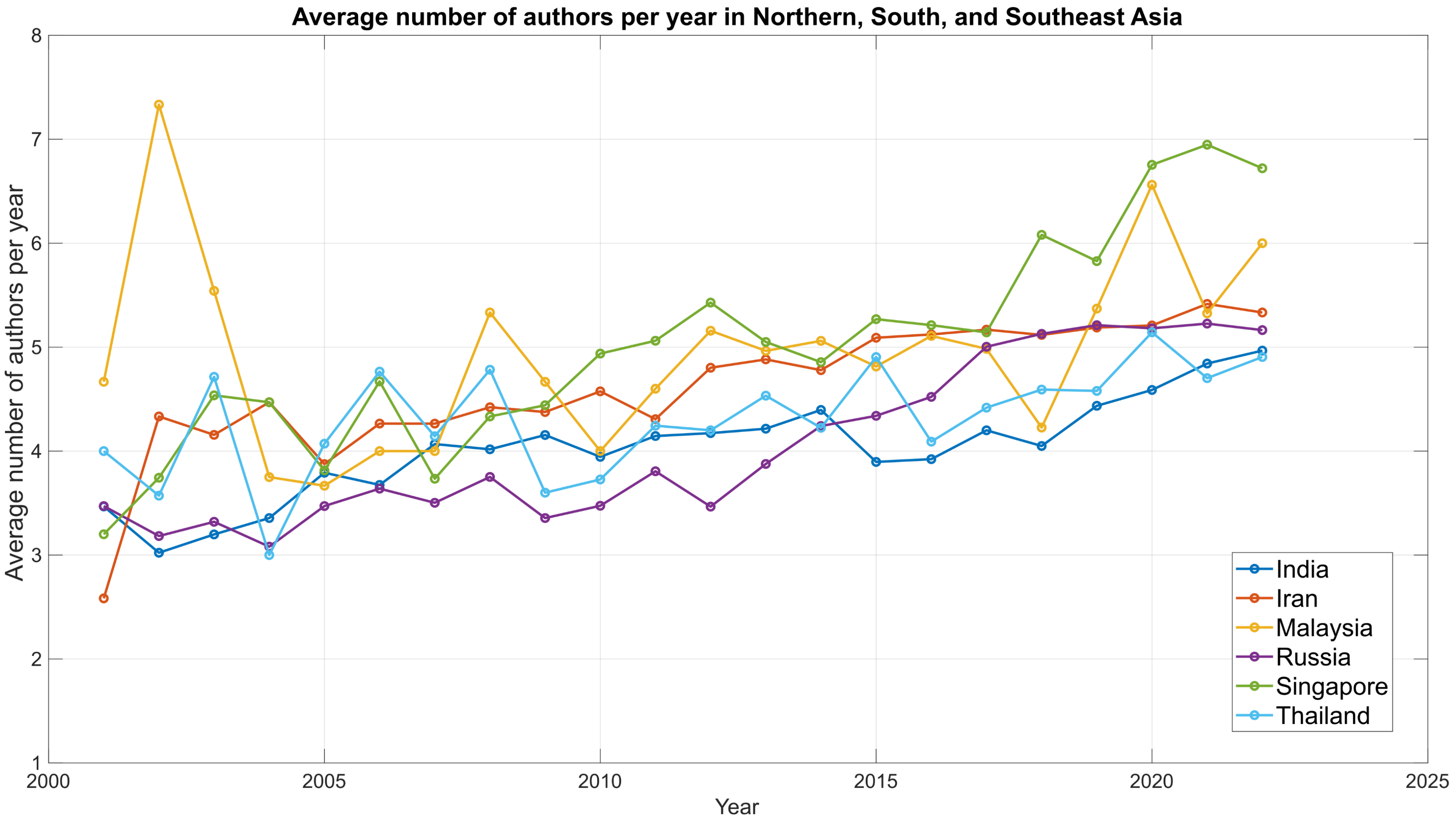
Average number of authors per year in Northern, South, and Southeast Asia
Average number of authors per year
Year
1
2
3
4
5
6
7
8
2000
2005
2010
2015
2020
2025
India
Iran
Malaysia
Russia
Singapore
Thailand

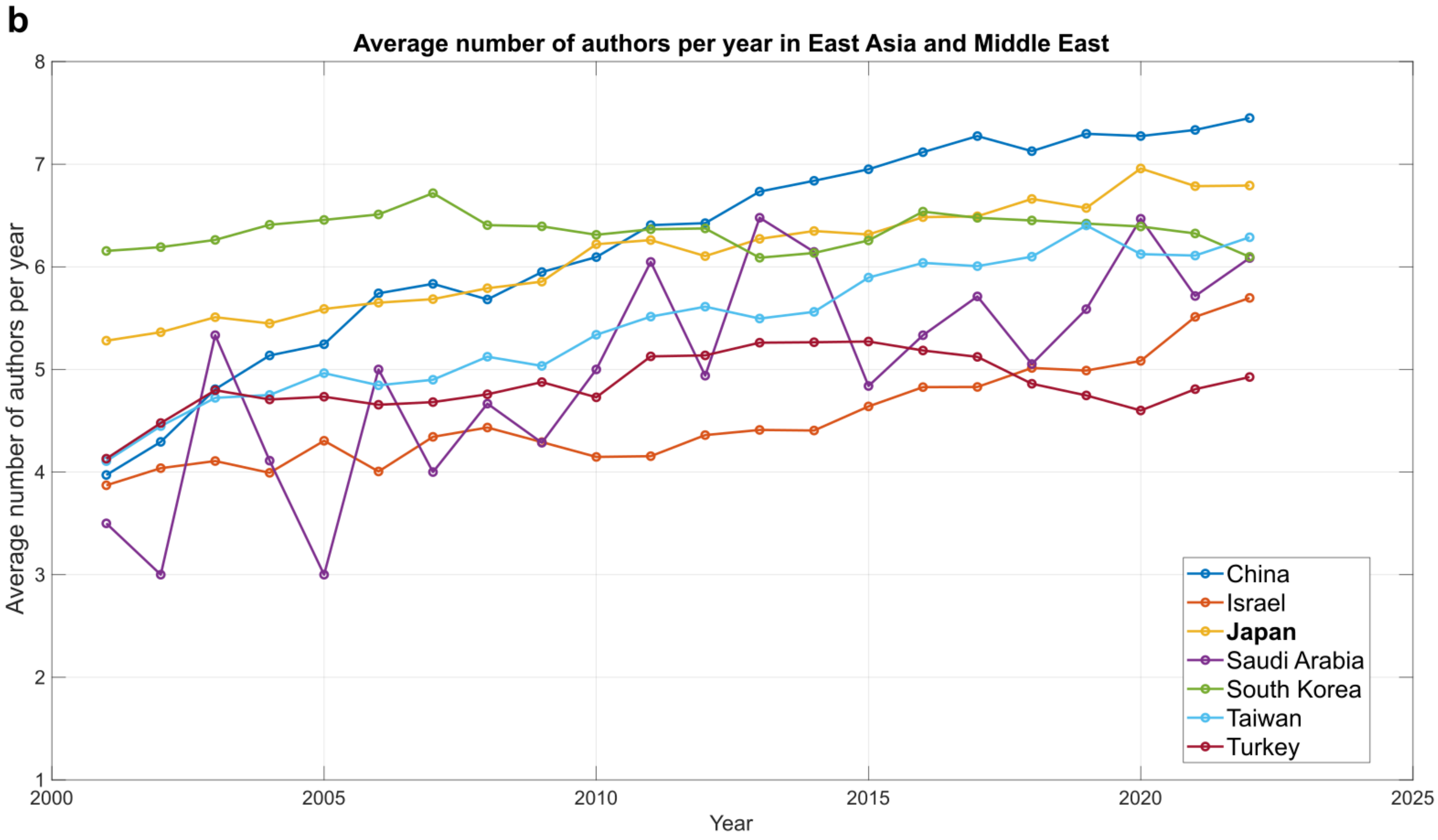


**Figure S3**. Temporal trend of authorship in the field of Neuroscience for the Asian target countries from 2001 to 2022. We divided the Asian countries into two separate figures following the regional classifications defined by WorldAtlas (www.worldatlas.com/), in order to avoid excessive overlapping between countries. **a**. Average number of authors per year in Northern, South and Southeast Asia. **b**. Average number of authors per year in East Asia and Middle East.

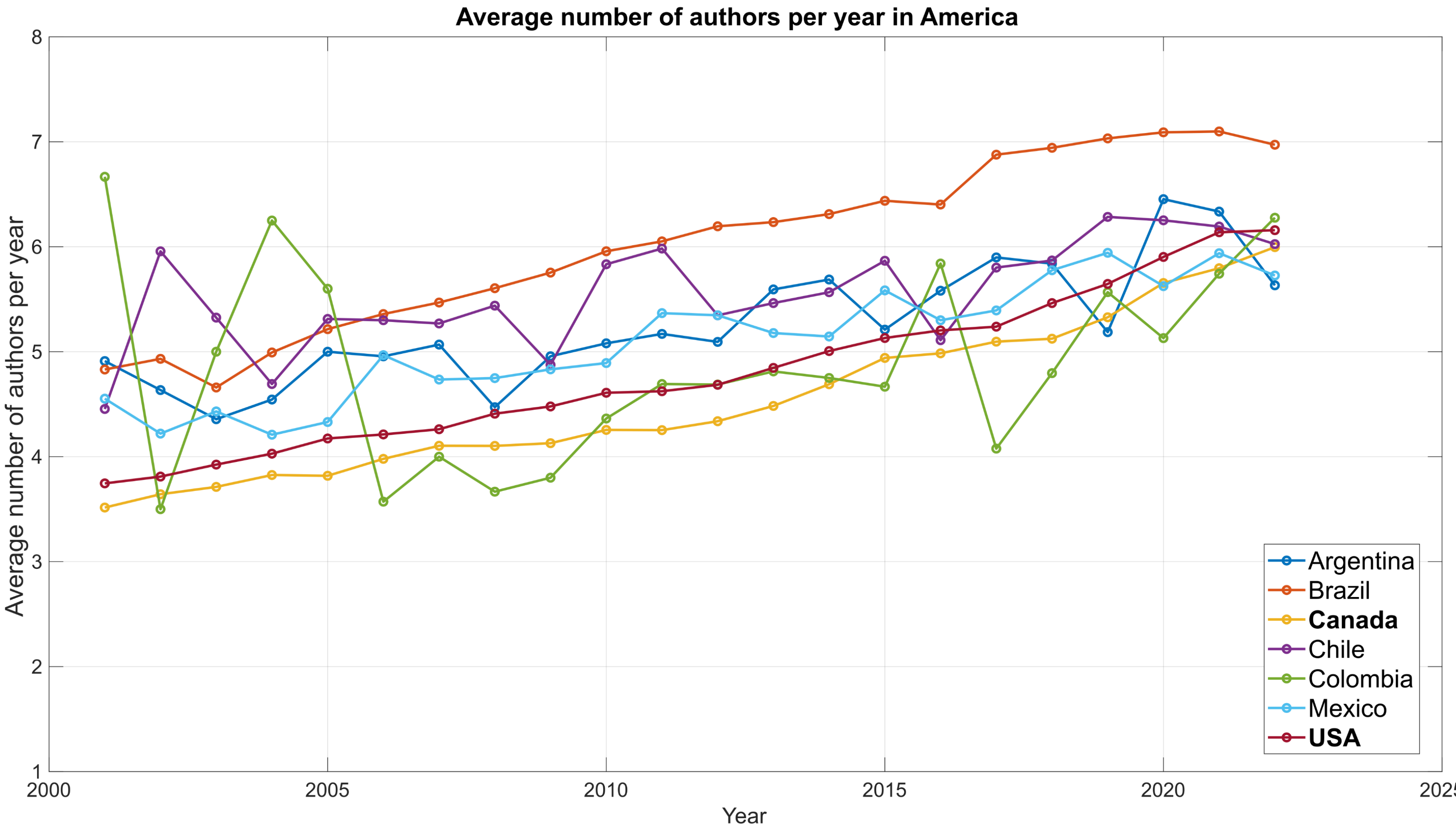


**Figure S4**. Temporal trend of authorship in the field of Neuroscience for the American target countries from 2001 to 2022.

**a**

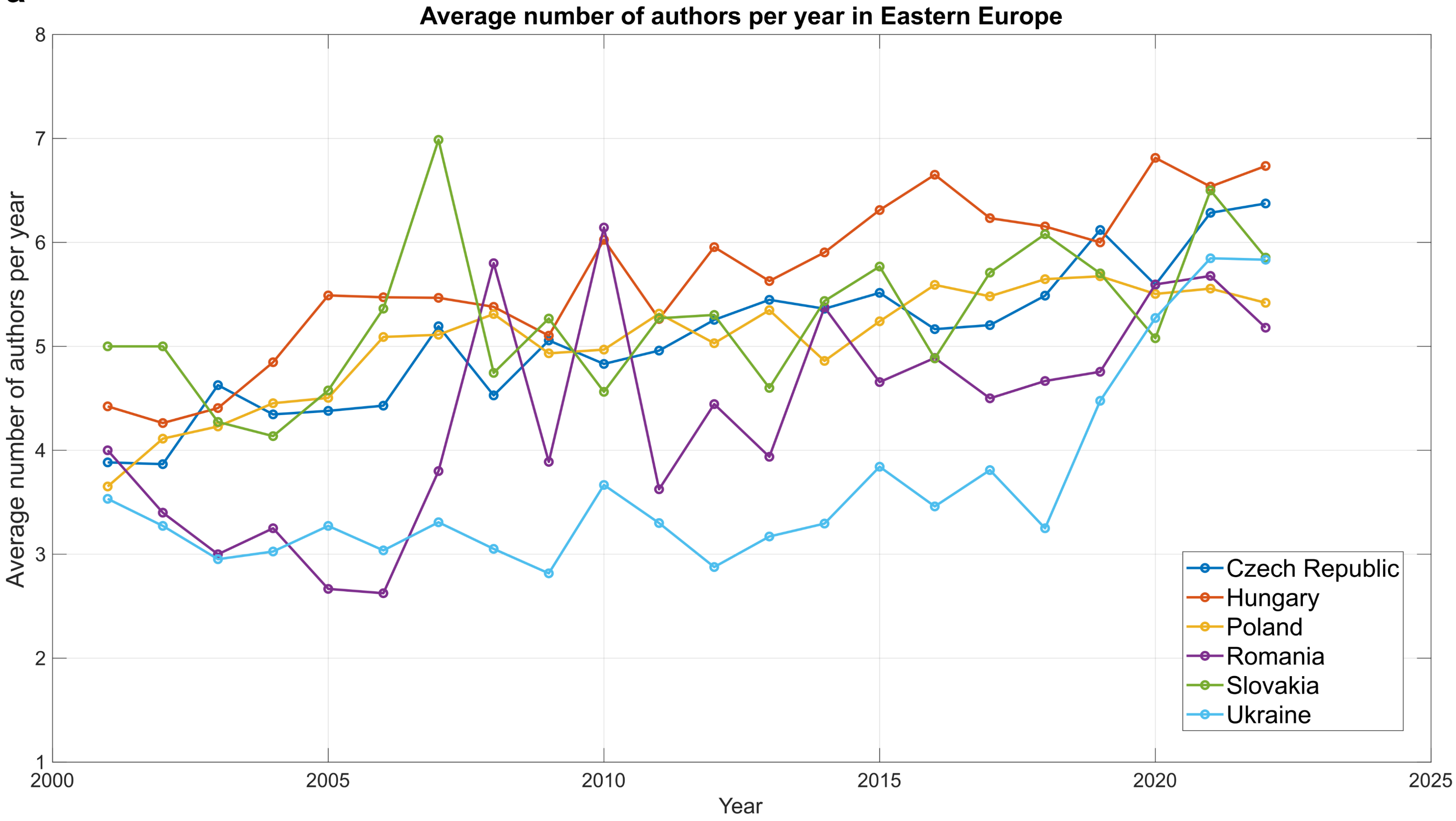
Average number of authors per year in Eastern Europe
Average number of authors per year
Year
8
7
6
5
4
3
2
1
2000
2005
2010
2015
2020
2025
Czech Republic
Hungary
Poland
Romania
Slovakia
Ukraine

b

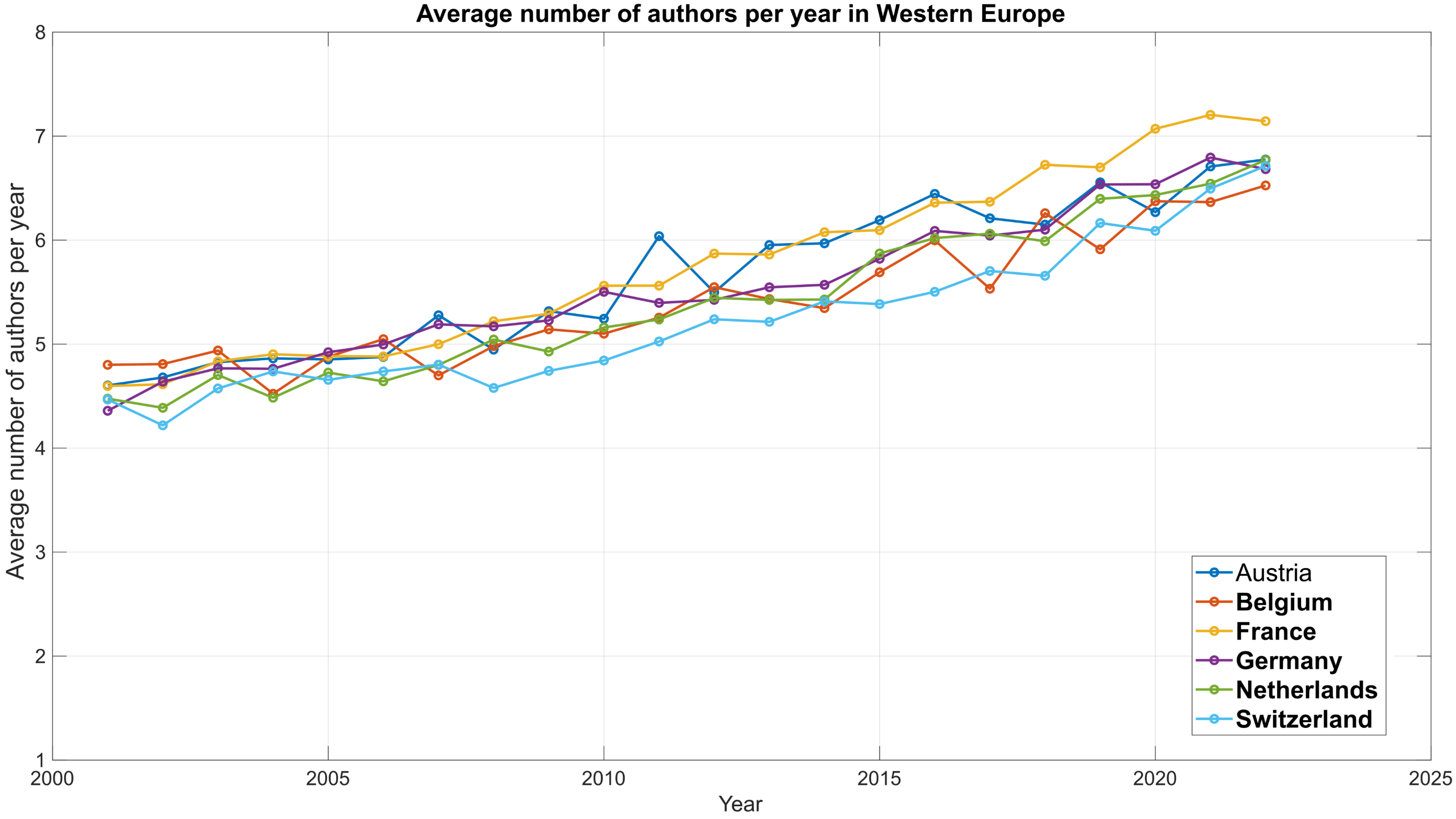
Average number of authors per year in Western Europe
Average number of authors per year
Year
Austria
Belgium
France
Germany
Netherlands
Switzerland
8
7
6
5
4
3
2
1
2000
2005
2010
2015
2020
2025

c

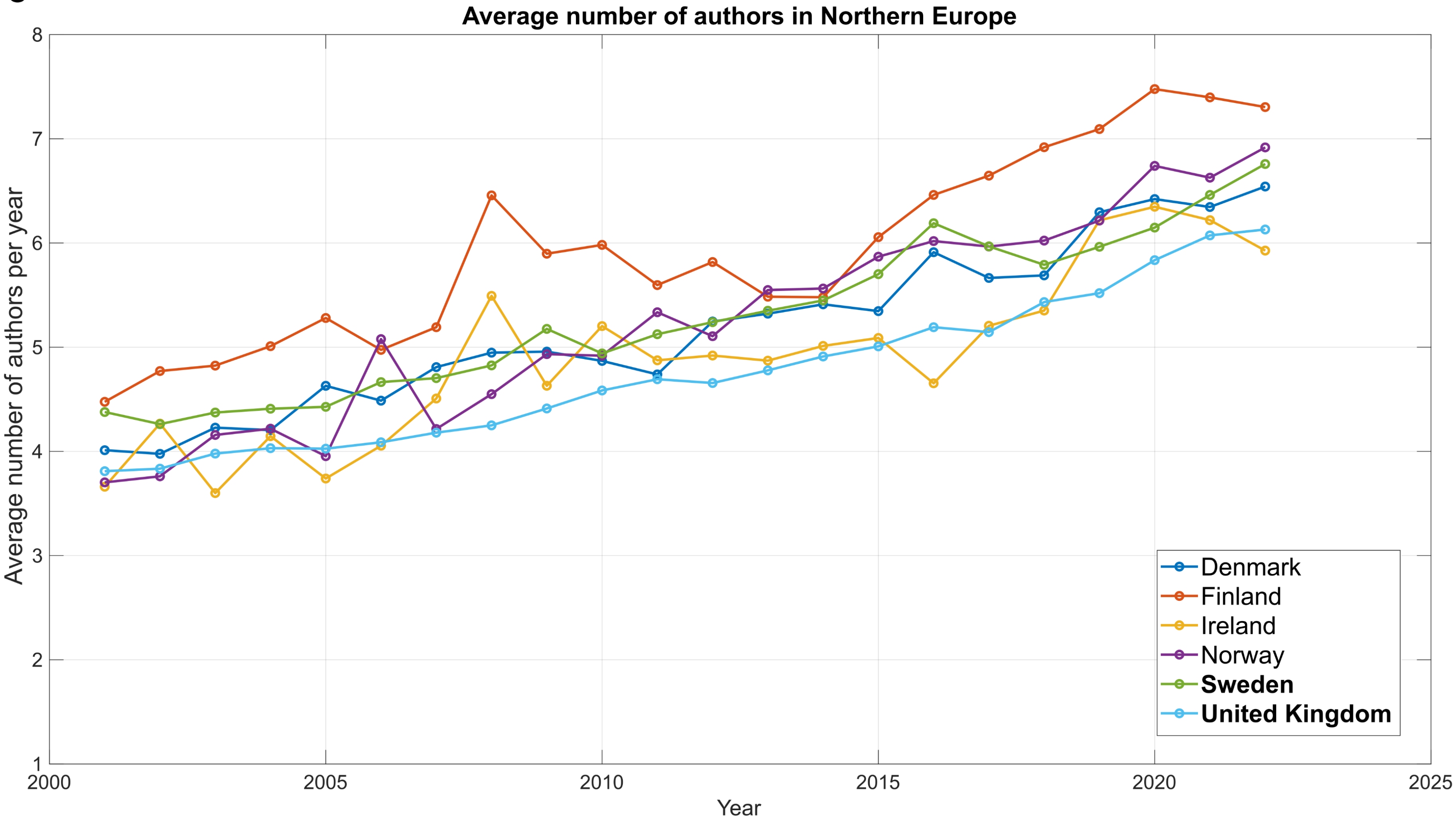

Average number of authors in Northern Europe
Average number of authors per year
Year
1
2
3
4
5
6
7
8
2000
2005
2010
2015
2020
2025
Denmark
Finland
Ireland
Norway
Sweden
United Kingdom

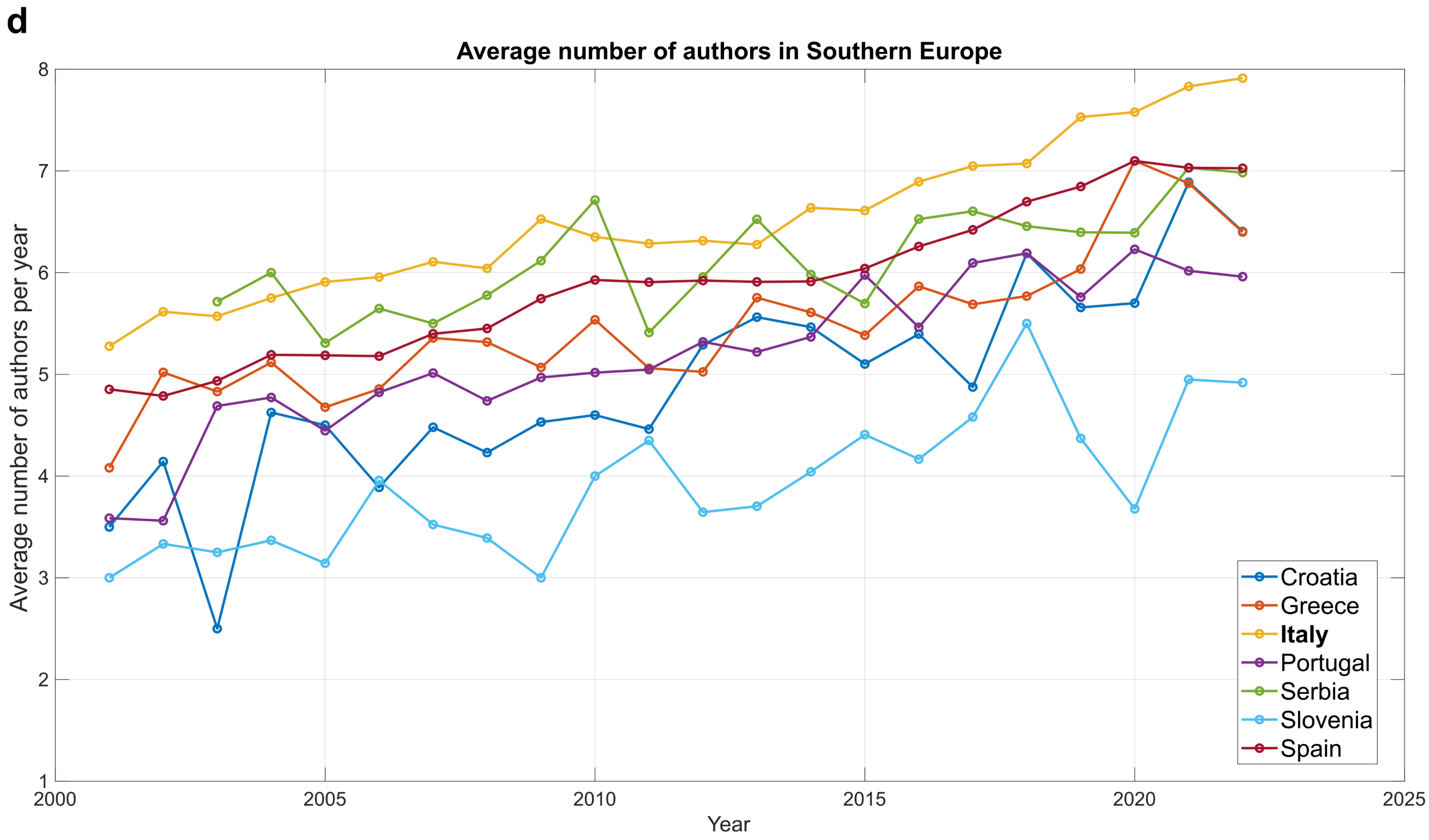


**Figure S5**. Temporal trend of authorship in the field of Neuroscience for the European target countries from 2001 to 2022. We divided the European countries into four separate figures to avoid excessive overlapping between countries. The division follows the regional classifications of Eastern, Western, Northern, and Southern Europe, as defined by the United Nations Geoscheme for Europe. **a**. Average number of authors per year in Eastern Europe. **b**. Average number of authors per year in Western Europe. **c**. Average number of authors per year in Northern Europe. **d**. Average number of authors per year in Southern Europe.

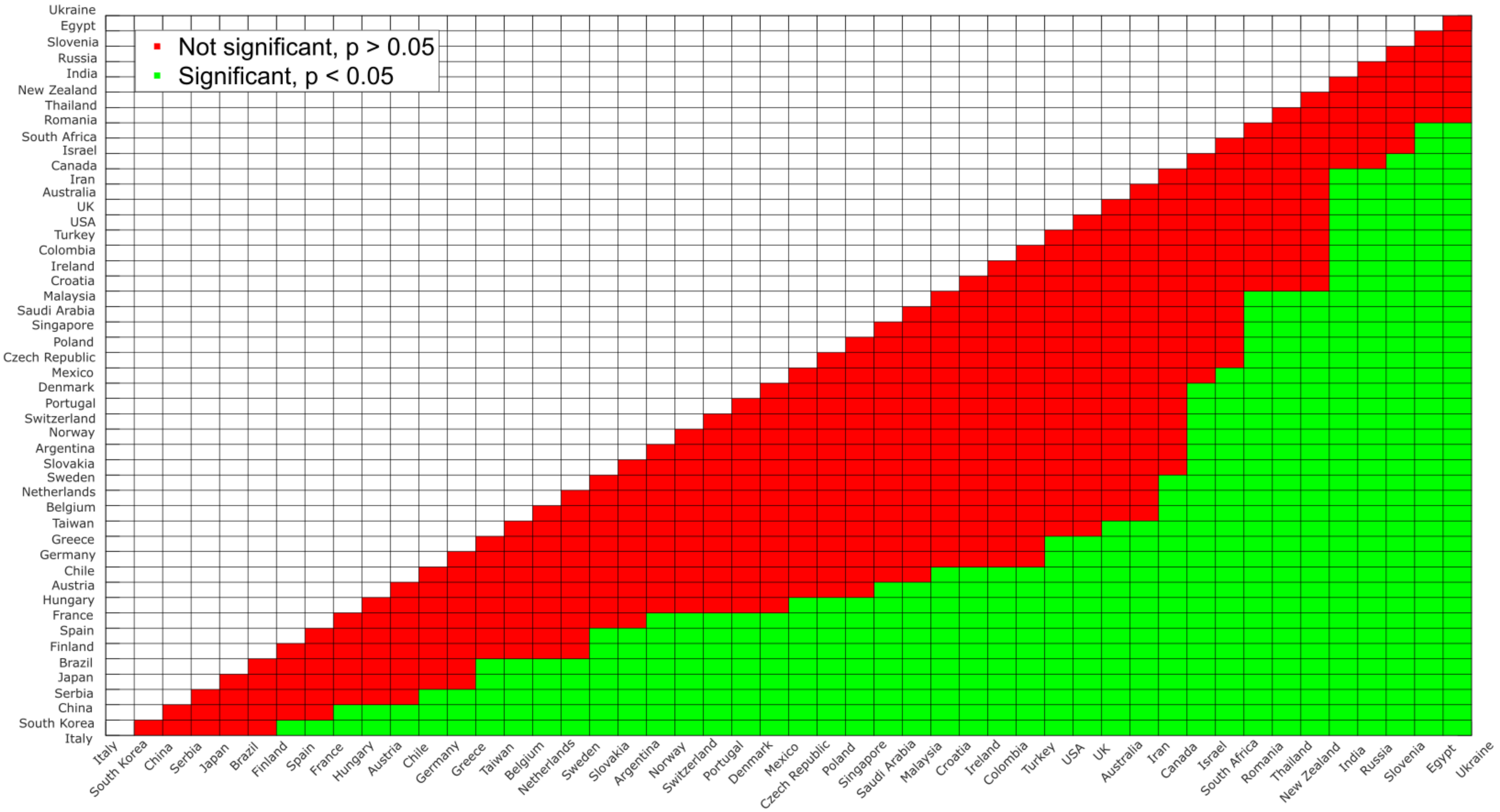


**Figure S6**. Country-by-Country Newman-Keuls post-hoc comparisons.

| | Country | Number of publications | Average of authors (2001-2022) | Average of authors (2001-2011) | Average of authors (2012-2022) | Δ Delta |
|---|---|---|---|---|---|---|
| 1 | **Italy** | 39212 | 6.50 (0.74) | 5.94 (0.37) | 7.06 (0.58) | 1.12 |
| 2 | South Korea | 13724 | 6.35 (0.15) | 6.38 (0.15) | 6.32 (0.15) | -0.06 |
| 3 | China | 69774 | 6.22 (1.03) | 5.37 (0.76) | 7.07 (0.30) | 1.70 |
| 4 | Serbia | 845 | 6.13 (0,51) | 5.79 (0.43) | 6.41 (0.40) | 0.62 |
| **5** | **Japan** | 46779 | 6.11 (0.51) | 5.69 (0.31) | 6.52 (0.25) | 0.83 |
| 6 | Brazil | 17737 | 6.01 (0.80) | 5.34 (0.46) | 6,69 (0.36) | 1.35 |
| 7 | Finland | 4947 | 5.93 (0.92) | 5,31 (0.60) | 6.55 (0.75) | 1.24 |
| 8 | Spain | 19612 | 5.89 (0.73) | 5.32 (0,40) | 6.46 (0.48) | 1.24 |
| **9** | **France** | 28939 | 5.76 (0.85) | 5.03 (0.33) | 6.49 (0.50) | 1.46 |
| 10 | Hungary | 3838 | 5.68 (0.75) | 5.10 (0.55) | 6.26 (0.38) | 1.16 |
| 11 | Austria | 5119 | 5.64 (0.72) | 5.04 (0.40) | 6.24 (0.36) | 1.20 |
| 12 | Chile | 1669 | 5.55 (0.50) | 5.31 (0.49) | 5.79 (0.38) | 0.48 |
| 13 | **Germany** | 55005 | 5.54 (0.69) | 4.99 (0.34) | 6.10 (0.48) | 1.11 |
| 14 | Greece | 2918 | 5.47 (0.70) | 4.99 (0.39) | 5.95 (0.61) | 0.96 |
| 15 | Taiwan | 7178 | 5.43 (0.64) | 4.88 (0.39) | 5.96 (0.29) | 1.08 |
| 16 | **Belgium** | 8584 | 5.42 (0.59) | 4.92 (0.21) | 5.90 (0.42) | 0.98 |
| 17 | **Netherlands** | 19674 | 5.40 (0.74) | 4.78 (0.28) | 6.03 (0.46) | 1.25 |
| 18 | **Sweden** | 10511 | 5.28 (0.74) | 4,66 (0.32) | 5.91 (0.46) | 1.25 |
| 19 | Slovakia | 1062 | 5.27 (0.70) | 5.01 (0.77) | 5.53 (0.54) | 0.52 |
| 20 | Argentina | 2992 | 5.25 (0.56) | 4.83 (0.27) | 5.68 (0.43) | 0.85 |
| 21 | Norway | 3959 | 5.24 (0.98) | 4.43 (0.55) | 6.05 (0.54) | 1.62 |
| 22 | **Switzerland** | 11736 | 5.22 (0.68) | 4.67 (0.21) | 5.77 (0.51) | 1.10 |
| 23 | Portugal | 3390 | 5.19 (0.75) | 4.60 (0.54) | 5.78 (0.37) | 1.18 |
| 24 | Denmark | 7271 | 5.18 (0.79) | 4.53 (0.37) | 5.83 (0.48) | 1.30 |
| 25 | Mexico | 4363 | 5.10 (0.54) | 4,66 (0.35) | 5.54 (0.28) | 0.88 |
| 26 | Czechia | 4355 | 5.08 (0.69) | 4,55 (0.43) | 5.61 (0.43) | 1.06 |
| 27 | Poland | 5996 | 5.04 (0.54) | 4.69 (0.54) | 5.39 (0.25) | 0.70 |

| | | | | | | |
|---|---|---|---|---|---|---|
| 28 | Singapore | 2499 | 5.01 (1.00) | 4.26 (0.57) | 5,75 (0,76) | 1.49 |
| 29 | Saudi Arabia | 778 | 5.01 (1.02) | 4.35 (0.96) | 5.66 (0.58) | 1.31 |
| 30 | Malaysia | 680 | 4.93 (0.90) | 4.60 (1.08) | 5.23 (0.61) | 0.63 |
| 31 | Croatia | 771 | 4.90 (0.99) | 4.13 (0.64) | 5.68 (0.59) | 1.55 |
| 32 | Ireland | 3258 | 4.90 (0.81) | 4.37 (0.62) | 5.43 (0.62) | 1.06 |
| 33 | Colombia | 481 | 4.88 (0.92) | 4.64 (1.11) | 5.12 (0.65) | 0.48 |
| 34 | Turkey | 7104 | 4.85 (0.28) | 4.69 (0.24) | 5.01 (0.23) | 0.32 |
| 35 | **USA** | 266333 | 4.80 (0.73) | 4.20 (0.30) | 5.40 (0.50) | 1.20 |
| 36 | **UK** | 53792 | 4.75 (0.71) | 4.17 (0.28) | 5.33 (0.50) | 1.16 |
| 37 | Australia | 25452 | 4.74 (0.89) | 4.01 (0.31) | 5.48 (0.63) | 1.47 |
| 38 | Iran | 6404 | 4.62 (0.63) | 4.14 (0.54) | 5.10 (0.20) | 0.96 |
| 39 | **Canada** | 40029 | 4.53 (0.73) | 3.93 (0.25) | 5.13 (0.52) | 1.20 |
| 40 | Israel | 8213 | 4.52 (0.49) | 4.15 (0.17) | 4.88 (0.43) | 0.73 |
| 41 | South Africa | 1178 | 4.36 (0.67) | 3.92 (0.45) | 4.80 (0.56) | 0.88 |
| 42 | Romania | 578 | 4.35 (1,02) | 3.83 (1,15) | 4.87 (0,53) | 1.04 |
| 43 | Thailand | 832 | 4.31 (0.52) | 4.05 (0.56) | 4.57 (0.32) | 0.52 |
| 44 | New Zealand | 3092 | 4.11 (0.65) | 3.63 (0.30) | 4.59 (0.55) | 0.96 |
| 45 | Russia | 4669 | 4.06 (0.76) | 3.45 (0.22) | 4.66 (0.61) | 1.21 |
| 46 | India | 12784 | 4.02 (0.48) | 3.71 (0.39) | 4.33 (0.35) | 0.62 |
| 47 | Slovenia | 553 | 3.92 (0.67) | 3.48 (0.43) | 4.36 (0.59) | 0.88 |
| 48 | Egypt | 1353 | 3.81 (1.20) | 3.00 (1.14) | 4.62 (0.52) | 1.62 |
| 49 | Ukraine | 1209 | 3.65 (0.89) | 3.20 (0.25) | 4.10 (1.08) | 0.9 |

**Table S1.** Average number of authors, along with the standard deviation, across all journals in the field of neuroscience for the period from 2000 to 2022. The data is further divided into two sub-periods: 2001–2011 and 2012–2022. Additionally, the table includes the delta value, representing the change between these two sub-periods, for each country. In the table, the names of the countries that we previously published in our earlier study (Paul et al., 2024) are highlighted in bold.

**LIST OF THE JOURNALS**

**Selected Category: NEUROSCIENCES.**

**Year: 2022.**

**Selected Category Schema: WOS.**

**Total number: 306.**

**Copyright (c) 2024 Clarivate.**

- Acs chemical neuroscience
- Acta neurobiologiae experimentalis
- Acta neurologica belgica
- Acta neuropathologica
- Acta neuropathologica communications
- Acta neuropsychiatrica
- Actas espanolas de psiquiatria
- Acupuncture & electro-therapeutics research
- AIMS Neuroscience
- Alzheimer's & Dementia: Diagnosis, Assessment & Disease Monitoring
- Alzheimers & Dementia-Translational Research & Clinical Interventions
- Alzheimers research & therapy
- Annals of clinical and translational neurology
- Annals of neurology
- Annals of Neurosciences
- Annual review of neuroscience
- Annual review of vision science
- Archives italiennes de biologie
- Archives of Neuroscience
- Arquivos de neuro-psiquiatria
- Asn neuro
- Audiology and neuro-otology
- Autonomic neuroscience-basic & clinical
- Basic and Clinical Neuroscience

- Behavioral and brain functions
- Behavioral and brain sciences
- Behavioral neuroscience
- Behavioural brain research
- Behavioural pharmacology
- Biological cybernetics
- Biological psychiatry
- Biological psychiatry-cognitive neuroscience and neuroimaging
- Bipolar disorders
- Bmc neuroscience
- Brain
- Brain and behavior
- Brain and cognition
- Brain and language
- Brain behavior and evolution
- Brain behavior and immunity
- BRAIN-Broad Research in Artificial Intelligence and Neuroscience
- Brain Circulation
- Brain Communications
- Brain-Computer Interfaces
- Brain connectivity
- Brain impairment
- Brain injury
- Brain pathology
- Brain research
- Brain research bulletin
- Brain sciences
- Brain stimulation
- Brain structure & function
- Brain topography
- Cellular and molecular neurobiology

- Cephalalgia
- Cerebellum
- Cerebral cortex
- Ceska a slovenska neurologie a neurochirurgie
- Chemical senses
- Chemosensory perception
- Clinical autonomic research
- Clinical eeg and neuroscience
- Clinical neurophysiology
- Clinical Neurophysiology Practice
- Clinical psychopharmacology and neuroscience
- Clocks & Sleep
- Cns & neurological disorders-drug targets
- Cns neuroscience & therapeutics
- Cognitive affective & behavioral neuroscience
- Cognitive computation
- Cognitive neurodynamics
- Cognitive neuroscience
- Cognitive systems research
- Cortex
- Current alzheimer research
- Current Behavioral Neuroscience Reports
- Current Developmental Disorders Reports
- Current neurology and neuroscience reports
- Current neuropharmacology
- Current neurovascular research
- Current opinion in behavioral sciences
- Current opinion in neurobiology
- Current opinion in neurology
- Developmental cognitive neuroscience
- Developmental neurobiology

- Developmental neuroscience
- Dialogues in clinical neuroscience
- Egyptian Journal of Neurology Psychiatry and Neurosurgery
- Encephale-revue de psychiatrie clinique biologique et therapeutique
- Eneuro
- Eneurobiologia
- Epilepsia open
- European journal of neurology
- European journal of neuroscience
- European journal of pain
- European neurology
- European neuropsychopharmacology
- Experimental brain research
- Experimental neurobiology
- Experimental neurology
- Fluids and barriers of the cns
- Folia neuropathologica
- Frontiers in aging neuroscience
- Frontiers in behavioral neuroscience
- Frontiers in cellular neuroscience
- Frontiers in computational neuroscience
- Frontiers in human neuroscience
- Frontiers in integrative neuroscience
- Frontiers in molecular neuroscience
- Frontiers in neural circuits
- Frontiers in neuroanatomy
- Frontiers in neuroendocrinology
- Frontiers in neuroinformatics
- Frontiers in neurology
- Frontiers in neurorobotics
- Frontiers in neuroscience

- Frontiers in synaptic neuroscience
- Frontiers in systems neuroscience
- Gait & posture
- Genes brain and behavior
- Glia
- Hearing research
- Hippocampus
- Human brain mapping
- Human movement science
- IBRO Neuroscience Reports
- IBRO Reports
- Ideggyogyaszati szemle-clinical neuroscience
- Ieee transactions on cognitive and developmental systems
- Indian Journal of Neurotrauma
- International journal of developmental neuroscience
- International journal of neuropsychopharmacology
- International journal of neuroscience
- International journal of psychophysiology
- International Journal of Tryptophan Research
- Jaro-journal of the association for research in otolaryngology
- Journal of alzheimers disease
- Journal of Alzheimers Disease Reports
- Journal of cerebral blood flow and metabolism
- Journal of chemical neuroanatomy
- Journal of clinical neurophysiology
- Journal of clinical neuroscience
- Journal of Cognitive Enhancement
- Journal of cognitive neuroscience
- Journal of comparative neurology
- Journal of comparative physiology a-neuroethology sensory neural and behavioral physiology

- Journal of computational neuroscience
- Journal of electromyography and kinesiology
- Journal of headache and pain
- Journal of Huntingtons Disease
- Journal of integrative neuroscience
- Journal of mathematical neuroscience
- Journal of molecular neuroscience
- Journal of motor behavior
- Journal of musculoskeletal & neuronal interactions
- Journal of neural engineering
- Journal of neural transmission
- Journal of neurochemistry
- Journal of neurodevelopmental disorders
- Journal of neuroendocrinology
- Journal of neuroengineering and rehabilitation
- Journal of neurogenetics
- Journal of neuroimmune pharmacology
- Journal of neuroimmunology
- Journal of neuroinflammation
- Journal of neurolinguistics
- Journal of neuromuscular diseases
- Journal of neuropathology and experimental neurology
- Journal of neurophysiology
- Journal of neuropsychiatry and clinical neurosciences
- Journal of neuroscience
- Journal of neuroscience methods
- Journal of neuroscience research
- Journal of neurotrauma
- Journal of neurovirology
- Journal of pain
- Journal of parkinsons disease

- Journal of physiology-london
- Journal of pineal research
- Journal of psychiatry & neuroscience
- Journal of psychopharmacology
- Journal of psychophysiology
- Journal of sleep research
- Journal of stroke & cerebrovascular diseases
- Journal of the history of the neurosciences
- Journal of the international neuropsychological society
- Journal of the neurological sciences
- Journal of the peripheral nervous system
- Journal of vestibular research-equilibrium & orientation
- Learning & memory
- Metabolic brain disease
- Molecular and cellular neuroscience
- Molecular autism
- Molecular brain
- Molecular neurobiology
- Molecular neurodegeneration
- Molecular pain
- Molecular psychiatry
- Motor control
- Multiple sclerosis journal
- Muscle & nerve
- Nature Aging
- Nature and science of sleep
- Nature human behaviour
- Nature neuroscience
- Nature reviews neuroscience
- Network neuroscience
- Network-computation in neural systems

- Neural computation
- Neural development
- Neural networks
- Neural plasticity
- Neural regeneration research
- Neurobiology of aging
- Neurobiology of disease
- Neurobiology of Language
- Neurobiology of learning and memory
- Neurobiology of stress
- Neurochemical journal
- Neurochemical research
- Neurochemistry international
- Neurocirugia
- Neurodegenerative diseases
- Neuroendocrinology
- Neuroendocrinology letters
- Neurogastroenterology and motility
- Neuroimage
- Neuroimaging clinics of north america
- Neuroimmunomodulation
- Neuroinformatics
- Neurologic clinics
- Neurological research
- Neurological sciences
- Neurological sciences and neurophysiology
- Neurology Research International
- Neurology india
- Neurology-neuroimmunology & neuroinflammation
- Neuromolecular medicine
- Neuromuscular disorders

- Neuron
- Neuropathology
- Neuropathology and applied neurobiology
- Neuropeptides
- Neuropharmacology
- Neuropsychopharmacology Reports
- Neurophotonics
- Neurophysiologie clinique-clinical neurophysiology
- Neurophysiology
- Neuropsychobiology
- Neuropsychologia
- Neuropsychological rehabilitation
- Neuropsychology
- Neuropsychology review
- Neuropsychopharmacology
- Neuroreport
- Neuroscience
- Neuroscience and biobehavioral reviews
- Neuroscience bulletin
- Neuroscience Insights
- Neuroscience letters
- Neuroscience research
- Neuroscientist
- Neurosonology and Cerebral Hemodynamics
- Neurotherapeutics
- Neurotoxicity research
- Neurotoxicology
- Neurotoxicology and teratology
- Neurotrauma Reports
- Npj parkinsons disease
- Npj science of learning

- Nutritional neuroscience
- Pain
- Pain Reports
- Pharmacology biochemistry and behavior
- Progress in neuro-psychopharmacology & biological psychiatry
- Progress in neurobiology
- Progress in Neurology and Psychiatry
- Psychiatric genetics
- Psychiatry and clinical neurosciences
- Psychoneuroendocrinology
- Psychopharmacology
- Psychophysiology
- Purinergic signalling
- Restorative neurology and neuroscience
- Reviews in the neurosciences
- Seizure-european journal of epilepsy
- Sleep
- Sleep and biological rhythms
- Sleep medicine reviews
- Social cognitive and affective neuroscience
- Social neuroscience
- Somatosensory and motor research
- Stereotactic and functional neurosurgery
- Stress-the international journal on the biology of stress
- Synapse
- Timing & Time Perception
- Translational neurodegeneration
- Translational neuroscience
- Translational stroke research
- Trends in cognitive sciences
- Trends in neurosciences

- Trends in Neuroscience and Education
- Vision research
- Visual neuroscience
- Zhurnal vysshei nervnoi deyatelnosti imeni i p pavlova
